# Divide and conquer: Using RhizoVision Explorer to aggregate data from multiple root scans using image concatenation and statistical methods

**DOI:** 10.1101/2024.07.05.602287

**Authors:** Anand Seethepalli, Chanae Ottley, Joanne Childs, Kevin Cope, Aubrey K. Fine, John Lagergren, Colleen M. Iversen, Udaya Kalluri, Larry M. York

**Affiliations:** Biosciences Division, Oak Ridge National Laboratory, Oak Ridge, TN, USA; Electrical and Computer Engineering Department, North Carolina State University, Raleigh, NC, USA; Environmental Sciences Division, Oak Ridge National Laboratory, Oak Ridge, TN, USA

## Abstract

Roots are important in agricultural and natural systems for determining plant productivity and soil carbon inputs. The collection of root samples from the field and their subsequent cleaning and scanning in a water-filled tray ranging in size from 5 to 20 cm, followed by digital image analysis has been commonly used since the 1990s for measuring root length, volume, area, and diameter. However, one common issue has been neglected. Sometimes, the amount of roots for a sample is too much to fit into a single scanned image, so the sample is divided among several scans. There is no standard method to aggregate the root measurements across the scans of the same sample. Here, we describe and validate two methods for standardizing measurements across multiple scans: image concatenation and statistical aggregation. Both methods rely on standardizing file naming conventions to identify scans that belong to the same sample. Image concatenation refers to combining digital images into a single larger image while maintaining the original resolution. We developed a Python script that identifies which images belong to the same sample and returns a single, larger concatenated image for every set of images in a directory. These concatenated images (combining up to 10 scans) and the original images were processed with RhizoVision Explorer, a free and open-source software developed for estimating root traits from images, with the same settings. An R script was developed that can identify the rows of data belonging to the same sample in RhizoVision Explorer data files and apply correct statistical methods such as summation, weighted average by length, and average to the appropriate measurement types to return a single data row for each sample. These two methods were compared using example images from switchgrass, poplar, and various tree and ericaceous shrub species from a northern peatland and the Arctic. Overall, the new methods accomplished the goal of standardizing measurement aggregation. Most root measurements were nearly identical except median diameter, which can not be accurately computed by statistical aggregation. We believe the availability of these methods will be useful to the root biology community.

## Introduction

Plants are drivers of terrestrial ecosystem processes and an integral component of global biogeochemical cycles. Plant productivity relies on belowground root systems, where larger-diameter coarse roots provide structural support for aboveground biomass, and narrower-diameter fine roots facilitate the acquisition of water and essential nutrients from the soil, and interface with soil microbes (Freschet et al., 2021). Root system architecture, the timing, placement, and structure of root systems, is linked to agricultural yield through water and nutrient uptake (Lynch et al., 2023). The rhizosphere, or zone of soil under the direct influence of living roots, is a hot spot of microbial activity fueled by direct inputs of photosynthetically-fixed carbon released to the soil during root growth, exudation, and turnover (Smithwick et al., 2014; York et al., 2016). Highly diverse rhizosphere microbial communities regulate nutrient bioavailability, expand the effective volume of soil accessed by roots, and modify the physical and chemical properties of soil (Hinsinger et al., 2009; Lambers et al., 2009). In managed and natural ecosystems, root activity and turnover initiates positive feedback loops between microbial activity and soil physicochemical structure that can promote carbon accrual as soil organic matter (Dijkstra et al., 2021). Understanding how roots and microbes interact in the soil is crucial for advancing bioenergy systems based on plant feedstocks (York et al., 2022). While the assessment of root morphological traits using image analysis has provided valuable insights into root biology, current approaches suffer from methodological challenges that still need to be addressed.

Many methods exist to measure root traits from images, but one of the most common due to its versatility and accuracy is washing roots clean from soil or pots, spreading them on a clear tray in water, and using a desktop scanner with a transparency unit to capture high contrast images of the roots. Over the past 30 years, numerous image analysis methods have been developed to identify roots in an image and measure their length, among other traits, with the first report of a recognizable segmentation to length measurement algorithm by Smucker et al. (1987). Pan and Bolton (1991) were the first to publish using a scanner for capturing images of spread roots and an analysis program based on the segmentation-feature analysis method (rather than line-intersection). The commercial software WinRhizo^™^ was released in 1993 (Arsenault et al., 1995) along with the sales of adapted transparency/film scanners to capture high-contrast images. WinRhizo^™^ is based on the same general algorithms described by Smucker et al. (1987) to measure root length, diameters, areas, and volume, but with additions like more thresholding options and topology analysis (see Seethepalli et al., 2021 for further historical details). However, the developer of WinRhizo^™^ did not release much validation data, some of the algorithms used are not known (so they are not reproducible), and its accuracy in measuring total volume using its default method was questioned (Delory et al., 2017; Rose, 2017; Rose and Lobet, 2019). While other software have been available over time, none quite captured the needs of the root, plant, and soil research communities to provide an easy-to-use, accurate, and reliable method for measuring roots. RhizoVision Explorer was released in 2021, along with extensive ground truth imagery with copper wires and roots from diverse species, was shown to be accurate for all traits including volume, and has been widely adopted by researchers around the world with more than 10,000 downloads and 100 citations at the time of this publication (Seethepalli et al., 2021). Despite this success, root biologists still face many challenges where multiple types of image analysis software remain needed and useful, and several issues remain unresolved.

One common issue is that root samples are frequently too large (i.e., contain too many roots) to fit in single scans without overcrowding. Commonly used flatbed scanners with transparency units, the Epson Expression 12000XL and Epson Perfection v850 have tray sizes of 420 cm x 300 cm and 280 cm x 210 cm, respectively. DeLory and colleagues (2017) demonstrated that root length density on a tray should not exceed 1 cm root cm^-2^ of tray, because as root overlap exceeds this threshold, the software is no longer able to accurately distinguish individual roots because the root edges merge with the background of another root. In this case, the software measurements will be inaccurate by measuring fewer but larger diameter predicted objects. The total root length for each sample from many experiments greatly exceeds this threshold, so researchers commonly divide roots over multiple scans. In this case, researchers commonly analyze each subsample image and manually sum the root lengths for a particular sample. However, they may not attempt to retain other measured traits due to the additional difficulty of aggregating calculations that emphasize statistical modes. Software like RhizoVision Explorer often have more than 10 measured traits that are useful for statistical analysis to distinguish treatment effects and to better understand root function. Multivariate methods are becoming more common and highlight the increased statistical power that retaining all measured traits after subsample scanning would provide (Mattupalli et al., 2019). Therefore, the root biology community would benefit from a standardized method to aggregate root trait measurements from multiple scans. Here, we provide two.

## Materials and Methods

The overall goal was to provide root biology researchers methods to handle multiple subsample scans of roots to retain all possible measurements provided by root image analysis software. These methods have been validated by three different case examples of roots from different species and systems, scanned on different model Epson scanners in different labs by different researchers. The root images are available (doi.org/10.5281/zenodo.12667583), the Python script for concatenation is available (doi.org/10.5281/zenodo.12668073), and the R script for statistical aggregation is available (doi.org/10.5281/zenodo.12668177). An updated protocol for scanning, including recommendations for handling multiple scans is available (doi.org/10.5281/zenodo.7825893).

### Description of validation root scans

Images were derived from three existing image sets that had multiple scans for the same sample all scanned on different scanners by different individuals using three species. In all cases, real examples of multiple scans for one complete sample were selected to give a range of subscans per sample from 1 to 10.

#### Roots Growing in Northern Organic Soils

*Peatland*. Images of fine roots sampled from dominant vascular plants in an ombrotrophic bog, including trees *Picea mariana* (Mill.) B.S.P. and *Larix laricina* (Du Roi), as well as ericaceous shrubs and sparse sedges in the *Carex* and *Eriophorum* genera, were generated as a part of the Spruce and Peatland Responses Under Changing Environments (SPRUCE) experiment (Hanson et al., 2017). The SPRUCE experiment is located in the ombrotrophic S1 peat bog at the Marcell Experimental Forest in Northern Minnesota, USA (47°30.4760′N; 93°27.1620′W; 418 m above mean sea level). SPRUCE encompasses twelve large enclosures (7-m tall by 12.8 m in diameter) that are subjected to a range of above- and below-ground warming (from +0 to +9 °C) and elevated carbon dioxide concentrations (∼900 ppm). Root ingrowth core methodology (e.g., (Malhotra *et* al., 2020) was used to collect newly-produced roots from each experimental enclosure over the course of a growing season (in this case, June to October 2017). Mesh cores filled with sterile, root-free peat were inserted into a premade hole, left in place for five months, then collected. Cores were frozen at -20 °C until processing.After thawing, fine roots were carefully removed from 10-cm depth increments using tweezers then cleaned with distilled water and small paint brushes. Cleaned fine roots were floated in distilled water in 10 cm by 15 cm trays and scanned using an Epson V700 Photo Scanner (Model J221A; Epson America Inc., Los Alamitos, CA, USA) with a transparency unit at 1400 dpi to resolve the pixels associated with narrow-diameter ericaceous shrub roots.

##### Arctic tundra

Roots of tundra graminoids were sampled in 2018 and 2021 in experiments associated with nitrogen or carbon isotopic labeling of monoculture tundra plants exposed to Zero Power Warming manipulations (Lewin et al., 2017) on the wet, polygonal tundra of the Barrow Environmental Observatory in Utqiaġvik, Alaska, USA. Fine roots and rhizomes of target plant species in selected monocultures were sampled from the surface organic layer using a tundra knife to cut a 9 cm by 9 cm square, and from deeper mineral soils to the frozen soil boundary using a 5-cm inner diameter soil corer. Soils were frozen until processing. After thawing, fine roots were carefully removed from organic or mineral soils using tweezers and cleaned with distilled water and small paint brushes. Cleaned fine roots were floated in distilled water in 10 cm by 15 cm trays and scanned using an Epson V800 Photo Scanner (Epson America Inc., Los Alamitos, CA, USA) with a transparency unit at 1400 dpi. Scans were either associated with the tundra grass *Arctagrostis latifolia* or tundra sedge *Carex aquatilis*.

For simplicity, all these samples are referred to as Peatland.

#### Populus

Hybrid aspen (*Populus tremula* x *alba*) root images were generated as part of a whole plant phenotypic characterization experiment. The root imaging dataset was collected from 18 plants consisting of four transgenic lines (*Aux/IAA* RNAi transgenics in auxin signaling pathway) and two control lines with three replicates each. Briefly, roots were collected from six-month-old, greenhouse-grown plants cultivated in Farfard 3B potting soil (Fafard Sungro #3B) under ambient conditions as described previously (Payyavula et al., 2022; Andrews et al., 2024). Root systems were gently washed to removesoil, separated into scannable portions, blotted to remove moisture, and stored in envelopes at 4 °C for up to one week before scanning. To obtain scanned images, root portions were floated in water-filled scanning trays, separated as needed with a paintbrush or tweezers to allow for a clean scan, and then scanned at 1200 DPI on an Epson Perfection V850 scanner with a transparency unit.

#### Switchgrass

Switchgrass (*Panicum vigarum*) root images derived from an experiment comparing two ecotypes under factorial combinations of nitrogen and water treatments (Griffiths et al., 2022). Plants were grown from clones in tall mesocosms in a sand, perlite, and vermiculite mixture, harvested at 115 days after sowing, and the roots were washed over a sieve. Roots were spread out in a tray filled with water and scanned on an Epson Perfection 12000XL scanner with a transparency unit at 600 DPI and saved as JPG with 95% quality (high quality).

### Image expectations and naming convention

RhizoVision Explorer can analyze most common image file types, including BMP, PNG, TIFF, and JPEG. The image concatenation script described below works for those as well. To automatically determine which subsample scans belong to which complete sample, naming files systematically is important. Both the image concatenation script and the statistical aggregation script assume the file names of multiple root subsample scans are named by appending the name of the root sample with a text string such as “_scan1”, “_scan2” or “_tray1”, “_tray2”. Subsamples belonging to the same complete sample are assumed to have the *identical* name to the left of the underscore (“_”) in the previous examples. We recommend the use of barcodes and barcode scanners to facilitate accurate file naming.

### Python scripts for image concatenation

The *group_images* and *concatenate_images* functions were created to process multiple scanned root images into a single image containing the samples to be analyzed. The functions use the open-source software packages NumPy (Numerical Python), pandas, and cv2 from OpenCV to read files and threshold, concatenate, and write images.

The *group_images* function allows the users to specify how they would like to group the images. The user must provide an image directory and either an identifier common to all the image names or a CSV file with two columns: a column for the image names and another for the group names. A data frame of names is generated after the image directory is searched for multiple image types: PNG, TIF, TIFF, JPG, JPEG, and BMP. If a common identifier is provided, the computer determines the base name or grouping factor based on the portion of the image’s name that appears before the common identifier. For example, images named ‘SpeciesA_scan1.png’, ‘SpeciesA_scan2.png’, and ‘SpeciesA_scan3.png’ will be grouped using ‘scan’ as the common identifier and ‘SpeciesA’ as the base name. If a CSV file is provided, the grouping factor should be specified in a column titled ‘basename.’

If a group contains multiple images, the process of creating a single image is performed with the *concatenate_images* function. Before concatenation, thresholding can be applied to grayscale images to convert the images to binary if the user provides an existing folder path to save the converted images and a threshold value. This function utilizes the ‘cv2.hconcat’ function to horizontally concatenate images that were grouped. If the images within a group have mismatched height sizes, the function pads the bottom of the shorter images with white pixels such that all the images have equal height. Otherwise, groups containing a single image will be saved without concatenation in the same user-specified folder as the concatenated images.

### Analysis in RhizoVision Explorer

The image sets used here were previously thresholded and the binary PNG images were used for image analysis in RhizoVision Explorer 2.0.3 (Seethepalli and York, 2021). The algorithms used in RhizoVision Explorer have been previously described and validated with ground truth images (Seethepalli et al., 2021). For each image set, the respective DPI was set in RhizoVision Explorer but otherwise settings were the same. The DPI used at scanning time for Peatland, Poplar, and Switchgrass were 1400, 1200, and 600, respectively, due to differences in scanning protocols across different labs and experiments. For all, the ‘Broken roots’ mode was used with a threshold of ‘200’ (arbitrary for a binary image), root pruning threshold was set to 5 pixels, and no diameter ranges were used. The independent subsample images and the concatenated images for each species were processed in different groups for a total of 6 output files, which are combined for further analysis during data analysis for this paper.

### R scripts for statistical aggregation

The *combinefeatures* function was created in R to aggregate features from multiple subscans for each sample. All the features except Average Diameter, Median Diameter, Maximum Diameter and Branching Frequency are aggregated by summing the features across all the scans for each sample. The Average Diameter, Median Diameter, and Branching Frequency are aggregated by calculating weighted mean across total root length of the sample. Since the Median Diameter cannot be found for the whole root sample, we calculated the mean of all Median Diameter values as the aggregation method. The weighted means method was determined to be mathematically equivalent to Branching Frequency aggregated by calculating the ratio of sum of branch points across all the subsample scans to the aggregated Total Root Length. The equation for the weighted means is:

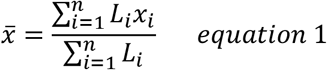

Where *x* is the respective root trait and *L* is the length of roots.

The Maximum Diameter is aggregated by finding the maximum value of all the Maximum Diameter values across all the subsample scans. While not included in the current study, this function will appropriately handle the additional traits provided by including additional diameter ranges in analysis, which are all traits aggregated through summation. The *combinefeatures* function relies on the *groupby* ability provided in *dplyr* from *tidyverse*. As shown in the provided R code, we assume researchers would split file names by underscores and at least one column (more are possible) are used in *groupby* to uniquely identify the complete sample such that subscans will be aggregated correctly. However, any method to identify which scans belong to the complete sample would work with minimal modification. Our methods provided here along with suggestions from our root scanning protocol () for naming conventions are meant to facilitate reproducible and convenient root scanning, image analysis, and data analysis.

### Statistical analysis

The *analysis*.*r* script was created in R to perform correlation analysis between the statistically aggregated features and features extracted from concatenated images for each sample. The script generates correlation plots and a table containing performance metrics for each feature. The correlation plots were made by ggplot() function. For each of these plots, the regression equation for each plant group (Peatland, Poplar or Switchgrass) was drawn using the stat_regline_equation() function. For calculating the performance metrics, the data was fit to linear models using the lm () function from stats R package for each imageset group. The script also depends on *tidyverse* R package for data manipulation, *ggpubr* R package for drawing regression equations and R-squared values on plots, *patchwork* R package to place multiple plots in a single figure grid and the *Metrics* package for calculating the performance metrics such as rmse() and mse().

## Results

To validate the two methods to combine root trait data from multiple subsample scans, we generated the same set of traits through both image concatenation and statistical aggregation as described above. We assume that image concatenation is the more accurate method because, after concatenation, roots are still analyzed using the same methods as for the individual images. In general, the results show that both methods give very similar results for all three image sets (Table 1, Figure 2).

**Table 1.**
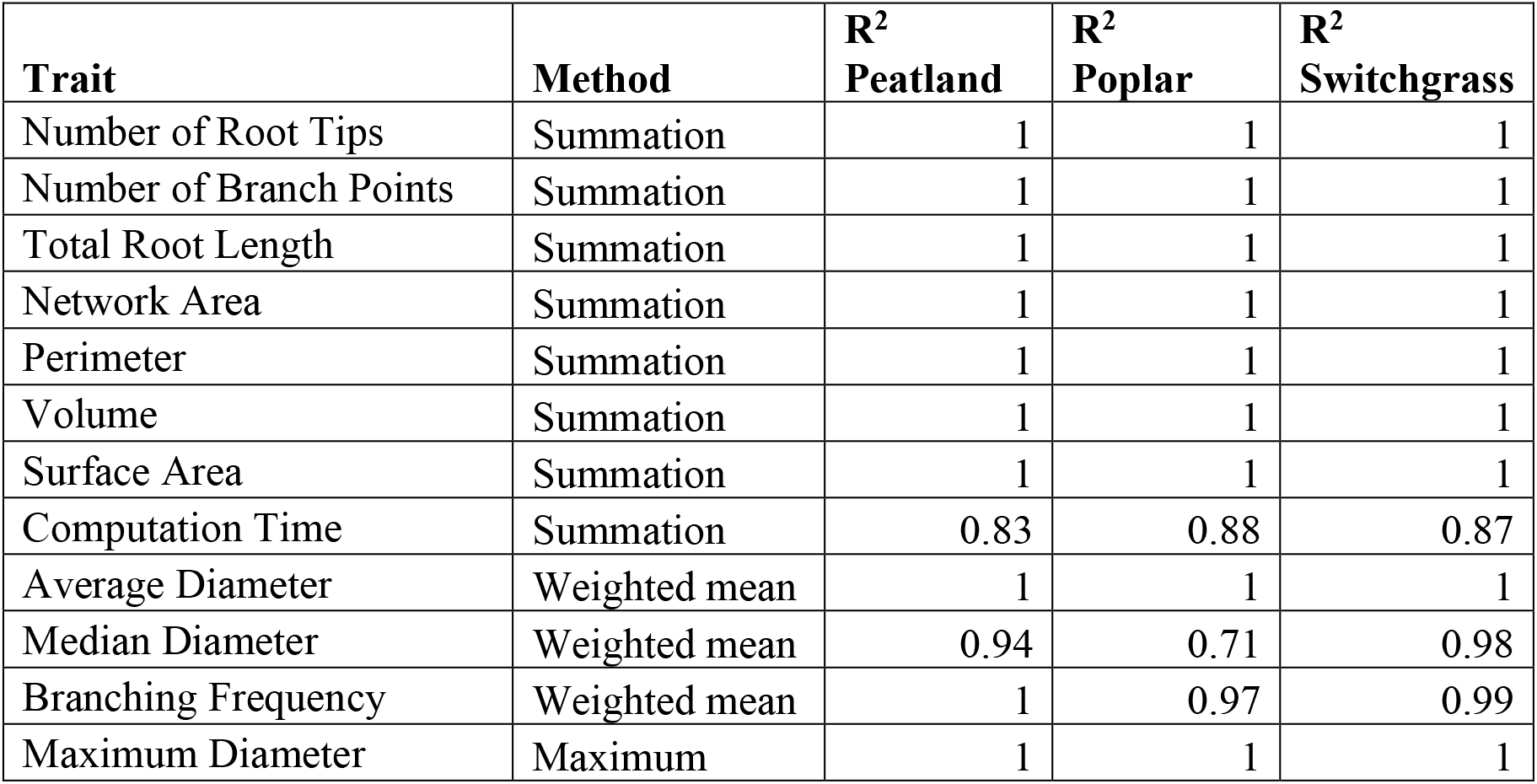
Summary of the methods to calculate each feature (or trait) using statistical aggregation across tabular data from RhizoVision Explorer and the R^2^ value of the regression of the values for each feature between concatenated image versus statistical aggregation.

| **Trait** | **Method** | **$R^2$**<br>**Peatland** | **$R^2$**<br>**Poplar** | **$R^2$**<br>**Switchgrass** |
| --- | --- | --- | --- | --- |
| Number of Root Tips | Summation | 1 | 1 | 1 |
| Number of Branch Points | Summation | 1 | 1 | 1 |
| Total Root Length | Summation | 1 | 1 | 1 |
| Network Area | Summation | 1 | 1 | 1 |
| Perimeter | Summation | 1 | 1 | 1 |
| Volume | Summation | 1 | 1 | 1 |
| Surface Area | Summation | 1 | 1 | 1 |
| Computation Time | Summation | 0.83 | 0.88 | 0.87 |
| Average Diameter | Weighted mean | 1 | 1 | 1 |
| Median Diameter | Weighted mean | 0.94 | 0.71 | 0.98 |
| Branching Frequency | Weighted mean | 1 | 0.97 | 0.99 |
| Maximum Diameter | Maximum | 1 | 1 | 1 |

**Figure 1.**
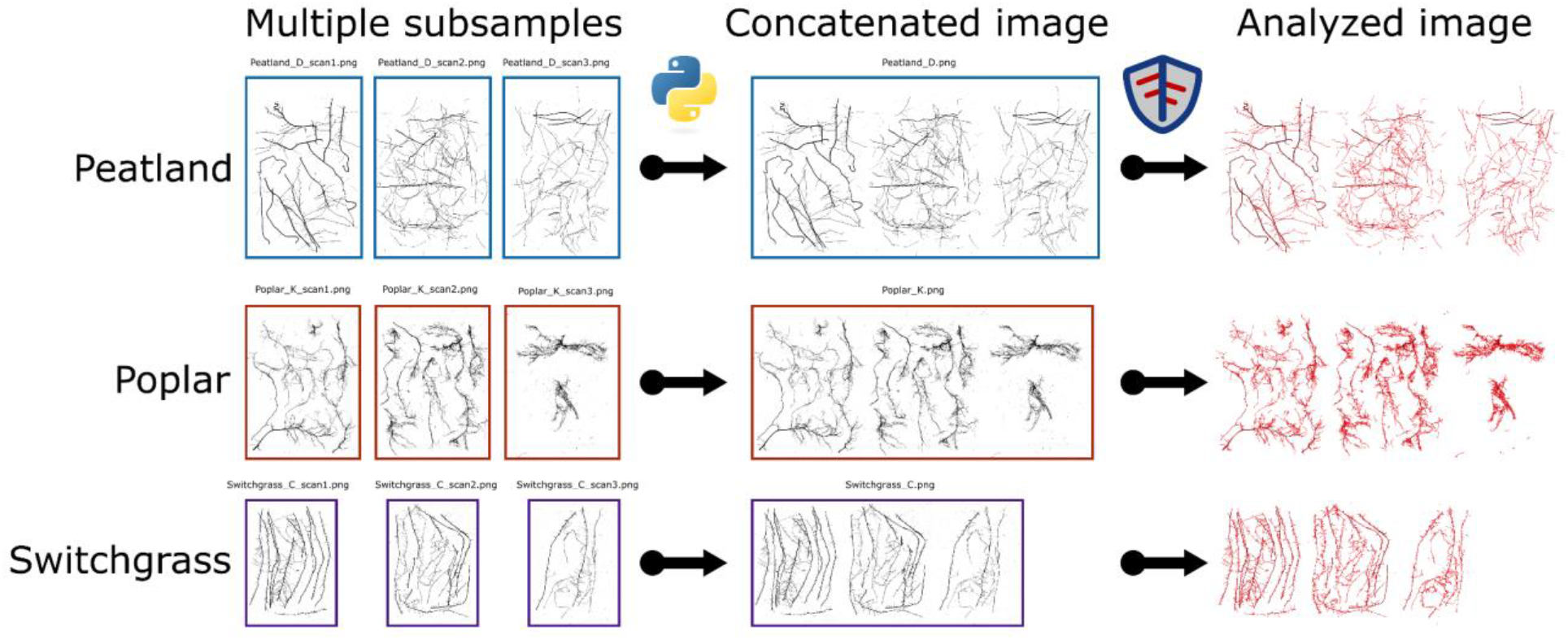
Multiple root subsample scans (left) are shown for root samples taken from a northern peatland, poplar, and switchgrass (top to bottom). For the image concatenation method, those subsamples were grouped horizontally into one large image using Python (center). These concatenated images were analyzed with RhizoVision Explorer (right) to produce root traits and compared to image analysis of the individual subsample scans with trait data combined through statistical aggregation.

**Figure 2.**
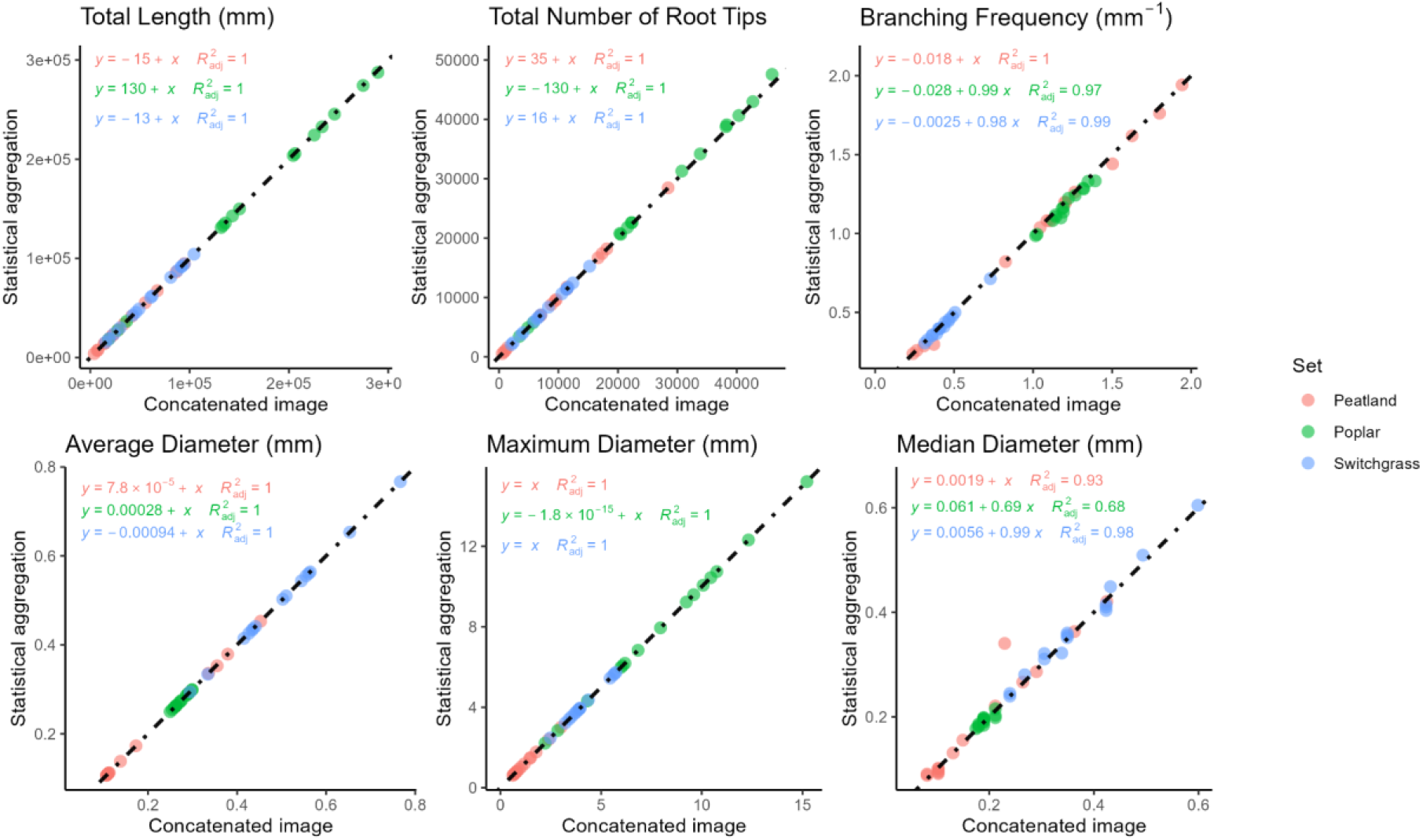
Six panels for representative types of root traits, including: Total length, Total Number of Root Tips, Branching Frequency, Average Diameter, Maximum Diameter, and Median Diameters. The dashed line depicts a 1:1 correspondence of values for each trait.

The root traits based on the statistical aggregation method of summation (Total Length, Total Number of Root Tips in Figure 2, Table 1) show nearly perfect agreement between the statistical aggregation method and the concatenation method with intercepts near 0, slopes near 1, and R^2^ near 1 for all image sets (Supplemental Table 1). For the weighted mean by length aggregations, Average Diameter and Branching Frequency showed nearly perfect agreement, but Median Diameter showed a lower R^2^, especially for the poplar image set (R^2^ = 0.71). The Maximum Diameter statistical aggregation method based on taking the maximum Maximum Diameter across the subsample scans had a nearly perfect agreement between the statistical aggregation method and the image concatenation method.

## Conclusions

These results indicate that both image concatenation and statistical aggregation provide similar results except for Median Diameter, which is influenced by the heterogeneity of diameters present across root classes and orders (Figure 2, Table 1). Average Diameter is more commonly used so this is not a major impediment to the statistical aggregation method. The concatenation and aggregation methods were implemented in Python and R, respectively, so are convenient for use in the plant science and ecology communities. We suspect the statistical aggregation method with an easy-to-use R function to combine features will be more widely used because more biologists and ecologists are likely familiar with R than Python. In the future, it may be possible to incorporate image concatenation into RhizoVision Explorer or other software, or manual image concatenation in software like GIMP would also work.

We believe this issue with multiple scans to be a common but disregarded issue for root image analysis and biology. Researchers may feel pressed to force as many roots as possible into a single scan instead, leading to possibly inaccurate measurements due to root overlap and occlusion as empirically determined (Delory et al., 2017). Or else, due to the perceived difficulties in data analysis, researchers may only aggregate a single trait across sub-scans manually, typically length. We hope these two methods, image concatenation and statistical aggregation, will allow more researchers to include more traits in the downstream analysis at a time when multivariate analysis and especially new approaches from machine learning, including deep learning, should be explored. Examples of multivariate data analysis for root traits include multiple regression (York and Lynch, 2015), principal component analysis for the root economic spectrum (Weigelt et al., 2021), classification using linear discriminate analysis (Mattupalli et al., 2019), and neural networks (Xu et al., 2022).

We note that other unresolved issues remain in root image analysis even with the relatively simple flatbed scanning of washed roots methods. These include the exclusion of root hairs from root identification that can drastically affect all measurements and better access to segmentation methods beyond the simple thresholding used in RhizoVision Explorer. Many recent papers have independently discovered and highlighted the synergy between use of RootPainter (Smith et al., 2022) for segmentation and RhizoVision Explorer for trait analysis (Alonso-Crespo et al., 2022; Bauer et al., 2022). Wider access to these technologies and AI would allow more routine use of minirhizotrons and rhizoboxes for *in situ* research with roots (Weihs et al., 2024). However, how to automatically identify the birth and death of roots as well as their longevity remains a challenge. The advancement of root image analysis allows new biological questions to be asked and answered, but possibly more importantly, the democratization of access to these tools is greatly expanding the number of people asking questions and we believe this greater access will drive the next major discoveries in root biology. These discoveries will be critical to mitigate climate change through soil carbon sequestration, understand natural ecosystems, and increase agricultural sustainability.

## Data Availability

The imageset is available at doi.org/10.5281/zenodo.12667583, the Python code for image concatenation is at doi.org/10.5281/zenodo.12668073, and the R code for statistical aggregation along with the figures and statistics presented here are available at (doi.org/10.5281/zenodo.12668177).

## Acknowledgements

This paper is dedicated to the memory of Joanne Childs, a pioneer in minirhizotron and scanned root image analysis. We would like to thank GEM for coordinating a summer fellowship for CO to work on this project and the Plant Microbe Interfaces SFA for providing funding for her fellowship and the time of AS, KC, JL, UK, and LY. This research was partially funded by the Genomic System Sciences Program, U.S. Department of Energy, Office of Science, Biological and Environmental Research, as part of the Plant-Microbe Interfaces Science Focus Area at the Oak Ridge National Laboratory (http://pmi.ornl.gov). This material is based upon work supported by the Center for Bioenergy Innovation (CBI), U.S. Department of Energy, Office of Science, Biological and Environmental Research Program under Award Number ERKP886. For JC and CI, the SPRUCE (Spruce and Peatland Responses Under Changing Environments) and Next-Generation Ecosystem Experiments in the Arctic (NGEE Arctic) projects are supported by the Biological and Environmental Research program in the U.S. Department of Energy’s Office of Science. We thank the UIC Science Native Corporation for their logistical support and for allowing us to conduct our science on the Barrow Environmental Observatory, lands of the Iñupiat since time immemorial. Oak Ridge National Laboratory is managed by UT-Battelle, LLC under Contract DE-AC05-00OR22725 with the U.S. Department of Energy (DOE).

## Notes

http://doi.org/10.5281/zenodo.12667583

http://doi.org/10.5281/zenodo.12668073

http://doi.org/10.5281/zenodo.12668177

### Competing Interest Statement

The authors have declared no competing interest.

